# Diversity in olfactory receptor repertoires is associated with dietary specialization in a genus of frugivorous bat

**DOI:** 10.1101/2020.12.31.424977

**Authors:** Laurel R. Yohe, Leith B. Leiser-Miller, Zofia A. Kaliszewska, Paul Donat, Sharlene E. Santana, Liliana M. Dávalos

**Author notes:** **Corresponding Author:** Laurel R. Yohe, 210 Whitney Ave., New Haven, CT 06511. **E-mail Addresses:** LRY LBM ZAK PD SES LMD.

## Abstract

Mammalian *olfactory receptors* (*OR*s) are a diverse family of genes encoding proteins that directly interact with environmental chemical cues. *OR*s evolve via gene duplication in a birth-death fashion, neofunctionalizing and pseudogenizing over time. Olfaction is a primary sense used for food detection in plant-visiting bats, but the relationship between dietary specialization and *OR* repertoires is unclear. Within neotropical Leaf-nosed bats (Phyllostomidae), many lineages are plant specialists, and some have a distinct *OR* repertoire compared to insectivorous species. Yet, whether specialization on particular plant genera is associated with the evolution of more specialized *OR* repertoires has never been tested. Using targeted sequence capture, we sequenced the *OR* repertoires of three sympatric species of short-tailed leaf-nosed bats (*Carollia*), which vary in their degree of specialization on the fruits of *Piper* plants. We characterized orthologous versus duplicated receptors among *Carollia* species, and identified orthologous receptors and associated paralogs to explore the diversity and redundancy of the receptor gene repertoire. The most dedicated *Piper* specialist, *Carollia castanea*, had lower *OR* diversity compared to the two more generalist species (*sowelli, perspicillata*), but we discovered a few unique sets of *OR*s within *C. castanea* with exceptional redundancy of similar gene duplicates. These unique receptors potentially enable *C. castanea* to detect *Piper* fruit odorants to an extent that the other species cannot. *C. perspicillata*, the species with the most generalist diet, had a larger diversity of functional receptors, suggesting the ability to detect a wider range of odorant molecules. The variation among *OR*s may be a factor in the coexistence of these sympatric species, facilitating the exploitation of different plant resources. Our study sheds light on how gene duplication plays a role in dietary adaptations and underlies patterns of ecological interactions between bats and plants.

**Impact Statement—though it asks for 3-4 sentences:** The sense of smell is essential to how many animals detect food, yet few studies have demonstrated how dietary evolution has shaped *olfactory receptor* genes, which encode proteins that bind to environmental scent cues, including food odorants. We compared the evolutionary history of olfactory receptor repertoires in three co-occurring neotropical bat species along a spectrum of dietary specialization on the fruits of *Piper* plants. We found the more generalist species possessed a more diverse olfactory receptor profile, potentially reflecting an ability to detect more diverse arrays of fruit scent compounds, while the specialist had a narrower profile that demonstrated more redundancy. By introducing creative approaches to measure diversity in large gene families and connecting diet specialization and molecular diversity, this study makes an unprecedented contribution to evolutionary biology.

## Introduction

The fitness of an animal is dependent on finding food, locating mates, and avoiding predation. The biochemical and cellular mechanisms underlying the sense of smell are excellent targets for natural selection because of their relevance to fitness and the ubiquity of chemosensation in animals (Hayden et al. 2010; Niimura 2012; Nikaido et al. 2013). To perceive a smell, an odorant molecule binds to its specialized olfactory receptor (*OR*) in a combinatorial fashion (Malnic et al. 1999; Nara et al. 2011; Kurian et al. 2020), precipitating a signaling cascade that ultimately transmits the odorant information to the brain. Each individual olfactory neuron expresses a unique *OR* allele; thus, its often interpreted that the larger the intact *OR* repertoire, the larger the combination of different odorants an organism can sense (Rodriguez 2013). This direct interaction with environmental signals suggests natural selection likely fine tunes *OR* binding motifs to optimally detect chemical cues relevant to fitness. However, deciphering the connection between *OR*s and the ecology of animals has proved challenging, because *OR*s evolve through paralogous duplication and the chemical cues necessary to elicit olfactory responses are complex (Yohe and Brand 2018).

*OR*s, as well as other many other chemosensory receptor genes, evolve in a birth-death manner, such that genes are constantly duplicating and pseudogenizing in tandem through time (Nei and Rooney 2005). This genetic mechanism of change has led to extraordinary diversity in chemoreceptor genes, making them among the largest and fastest-evolving protein-coding genes in the vertebrate genome (Niimura and Nei 2007; Nei et al. 2008; Niimura 2013; Yohe et al. 2020b). Variation in numbers of intact *OR* gene copies and *OR* pseudogenes can vary by orders of magnitude (Niimura et al. 2014). The fate of a gene duplicate includes one of several paths (Hahn 2009; Teufel et al. 2016; Yohe et al. 2019b). First, the duplicated gene may be completely redundant and not be expressed, and thus accumulate deleterious mutation that may render it a pseudogene (Eyun 2019). Second, one of the two copies may be released from purifying selection and accumulate new mutations that enable new function (Pegueroles et al. 2013). Third, the second copy may have a dosage effect, such that there is now increased the expression of the ancestral single copy (Loehlin and Carroll 2016), and fixation of the same copy of the gene is advantageous to fitness.

Measuring adaptation at the species level in large gene families has proven difficult, because of the challenges of simultaneously tracking both orthology versus paralogy and the rate of adaptive substitution (Hahn 2009; Han et al. 2009; Yohe et al. 2019b). Here we present a new approach to understanding the evolutionary history of *OR* gene duplicates among recently diverged species, in which we detect orthologous genes and associated paralogs from unrooted codon model gene trees and then measure diversity and evenness using metrics from community ecology. We apply this method to investigate *OR* diversity and evolution in three sympatric species of short-tailed leaf-nosed bats (*Carollia* spp.). The *Carollia* system is ideal for investigating a connection between ecological specialization and *OR* diversity for two reasons. First, all three species consume fruits of the genus *Piper*, but the degree of *Piper* specialization varies from *Carollia castanea* almost exclusively feeding on *Piper* fruits throughout the year, to *C. perspicillata* consuming only about 50% of *Piper* fruits and a variety of other plant genera from several families as well as nectar from flowers and insects occasionally, and *C. sowelli* falling somewhat in between the other two species (Fig. 1A; (Fleming 1991; Lopez and Vaughan 2007; Maynard et al. 2019). Second, behavioral assays have revealed that *Carollia* primarily use their sense of smell to initially locate fruiting patches and individual fruits, with echolocation being used at closer range to pinpoint the target fruit before grabbing it (Thies et al. 1998). *Carollia* also only seems to perform feeding attempts in the presence of scent cues from *Piper* fruit (Thies et al. 1998; Leiser-Miller et al. 2020).Thus, *Carollia*’s reliance on olfaction to locate *Piper* for fruits (and reciprocal reliance of *Piper* on chemical cues to attract *Carollia* for seed dispersal) makes it likely evolution has optimized the *OR* repertoires of each of these bat species for food detection. We predicted that, because *C. castanea* primarily needs to locate ripe *Piper* fruits, the bouquet of potential odorant ligands and therefore the diversity of respective receptors might be narrower than those of *C. perspicillata*, which needs to detect not just ligands from *Piper*, but also from the diversity of other plant foods it consumes. We apply our novel approach of using ecological metrics of diversity to measure diversity among orthologous and paralogous genes to investigate how evolution has shaped *OR* repertoires in the context of specialist and generalist diets.

**Figure 1.**
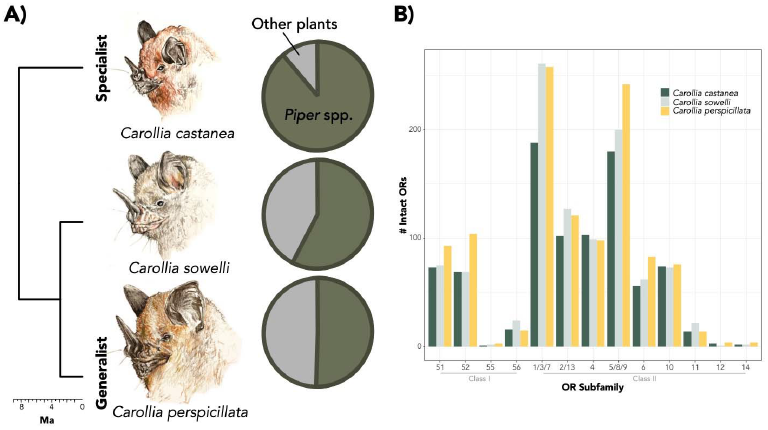
Target species of study that demonstrate a continuum of *Piper* specialization. (A) Proportion of *Piper* species found in diet of each *Carollia* species (based on Fleming (1991); Lopez and Vaughan (2007); and Maynard et al. (2019)). (B) Number of intact *olfactory receptor* (*OR*s) genes from sequence capture analysis within each subfamily. Illustrations by Christina M. Mauro.

## Methods

### Sampling and Sequencing

To test whether specialist and generalist species had distinct receptor profiles, we sequenced the *OR*s of three *Carollia* species using targeted sequence capture of probes designed from transcriptomic data. Samples were collected at La Selva Biological Station in Costa Rica during an August 2017 expedition. One male individual of each of the three *Carollia* species present at La Selva was captured on the evening August 4, 2017 at the same locality within the station (Table S1). Bats were trapped in mist nets and immediately placed in cloth bags prior to processing. Bats were euthanized using isoflurane and liver dissections were performed according to published video protocols (Yohe et al. 2019a). Bats and samples were processed in accordance with Stony Brook University Institutional Animal Care and Use Committee protocol #448712-3. Samples were collected with Costa Rica research permit #R-041-2017 and samples were exported from Costa Rica in alliance with country guidelines and imported following U.S. Center for Disease Control and U.S. Fish & Wildlife guidelines (USFW 3-177 2018NY2190224). For the targeted bait capture, probes were designed from a previously published analysis (Yohe et al. 2020a). Briefly, chemosensory receptors were identified in the transcriptomes of the main olfactory epithelium in twelve species of bats and probes were subsequently designed from the diversity of these receptor transcripts. DNA was extracted from flash-frozen liver tissue stored in RNA-later using the Qiagen QIAamp DNA Micro kit (Qiagen 56304). DNA quality was assessed using 260/280 ratios in a nanodrop, and DNA was quantified using a Qbit. DNA extractions were sent to Agilent where the chemoreceptor probes amplified *OR*s. Amplified targets were sequenced using Illumina HiSeq sequencing technology with 100bp paired-end reads by Arbor Biosciences (Ann Arbor, MI).

### Quality control and assembly

All sequence bait capture assemblies were performed using previously published methods optimized for large multigene families (Yohe et al. 2020a). Briefly, four lanes of raw paired end reads were generated for each species. Raw reads were trimmed using the bbduk.sh script in the BBTools genomic tools suite, in which regions with a quality score of less than 10 were trimmed. Using the bait designs as guides for assembling the raw reads, we implemented the reads_first.py in the HybPiper toolkit (Johnson et al. 2016). Each lane was assembled individually, and then resulting receptors were pooled, and duplicates were removed.

### Olfactory receptor annotation

In both the transcriptome assembly output and cleaned targeted bait capture output, contigs were run through the Olfactory Receptor Assigner v. 1.9.1, in which *OR*s were binned into respective subfamilies (Hayden et al. 2010). Pseudogenes were determined as open reading frames disrupted by a frameshift or premature stop codon mutation, or sequences less than 650bp. Pseudogenes were removed from the analysis.

### Alignment and gene tree inference

Each subfamily of intact receptors was aligned using transAlign (Bininda-Emonds 2005) option in Geneious v. 10.2. 3 (Kearse et al. 2012) with a MAFFT v. 7.388 (Katoh and Standley 2013) with the FFT-NS-2 algorithm for the protein alignment. The human adenosine A2b receptor, an ancestral G-protein-coupled receptor gene, was included in each alignment in order to root the gene trees (NM_000676.2), as suggested from previous publications on mammalian *OR*s (Niimura 2013). For model selection and tree inference, stop codons were removed. Model selection was performed on each alignment using ModelOMatic v. 1.01 (Whelan et al. 2015), in which 75 nucleotide, amino acid, and codon models were tested. Maximum likelihood tree inference was performed on each alignment with the estimated best-fit model using IQ-TREE v. 1.6.11 (Nguyen et al. 2015) with 1000 ultrafast bootstrap replicates.

### Orthogroup characterization

To characterize orthologous *OR* genes, as well as associated duplicates accumulated both prior to (out-paralogs) and after species divergence (in-paralogs), we used an unrooted phylogenetic assessment of the gene trees for each subfamily (Ballesteros and Hormiga 2016). For each gene tree, we used the UPhO.py script within UPhO implemented with Python v. 2.7.15 with the -iP flag to track in-paralogs and minimum number of species in an orthogroup to 1 (Ballesteros and Hormiga 2016).

### Receptor diversity and evenness

To quantify *OR* gene “diversity”, we used diversity indices commonly used in community ecology. The diversity of community composition is often assessed with species richness (number of species) and species abundances (number of individuals per species) at different sites within a community. These metrics are then used to calculate evenness (*J*’) across sites and community diversity (*e*.*g*. Shannon’s *H*’). Applying this framework, we considered each *OR* subfamily as a “community” and each gene orthogroup a “site” within the community. Instead of measuring abundance as number of individuals per species within a site, we measured number of genes (duplicates) per species within the orthogroup. Evenness reflects the relative abundance of each species represented within each orthogroup. The higher the similarity in abundance is across species, the higher the evenness. We can then calculate Shannon’s *H*’ and evenness *J*’ for total *OR* gene repertoires, as well as for each *OR* gene subfamily. Diversity indices were calculated using the diversityresult() function within the BiodiversityR v. 2.12.1 (Kindt 2016) in R. v. 4.0.2 (R Core Team 2020). To statistically compare diversity and evenness values among species, we performed a phylogenetically-corrected linear mixed effects model using the lme() function with the nlme v. 3.1.151 (Pinheiro et al. 2020) where the phylogenetic distance among species was measured from a correlation matrix estimated from a previously published phylogeny (Rojas et al. 2016).

## Results

### Olfactory receptor distribution

For each *Carollia* species, the number of intact *OR* genes are as follows: *C. castanea* with 881 OR genes, *C. sowelli* with 1017, and *C. perspicillata* with 1115 genes (Fig. 1B; Table S2). Figure 1B shows the abundance of ORs within each subfamily for each species. OR1/3/7 and OR5/8/9 show twice the abundance relative to other subfamilies for all species while subfamily OR55, OR12, and OR14 are represented in fewer paralogs relative to other subfamilies.

### Alignment and orthogroup inference

Alignments for each subfamily resulted in lengths ranging from 1065bp to 1242bp. For every alignment, codon models were the best fit models of evolution, though the state frequencies varied (Table S3). For all identified gene trees, a total of 1019 orthogroups were identified (Fig. 2). The number of orthogroups per subfamily are listed in Table S2. Alignments, gene trees, and orthogroup cluster lists are available in the supplement. Figure 2B indicates the abundance of receptors for each orthogroup for each OR subfamily, demonstrating how some orthogroups have higher abundances in some species versus others.

**Figure 2.**
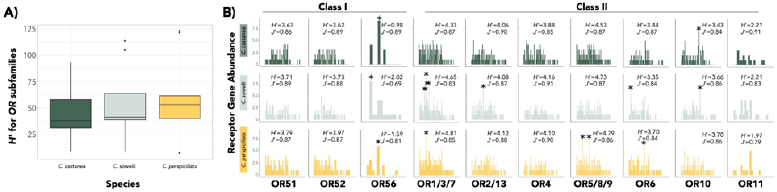
(A) Distribution of Shannon Diversity Indices (*H*’) for each *olfactory receptor* (*OR*) gene subfamily for each species. Phylogenetic linear mixed models result in *C. perspicillata* being significantly more diverse. The y-axis is on the exponential scale to emphasize differences (B) Abundance profiles for each species (*C. castanea*: evergreen; *C. sowelli*: mint green; *C. perspicillata*: gold) for each *OR* gene subfamily. Each bar denotes a unique orthogroup and the same orthogroup index is consistent across species for comparison. The Shannon Diversity Index (*H*’) and the evenness (*J*’) is presented for each “community” of genes. OR55, OR12, and OR14, which had only a few genes in each species, are not shown. * indicates orthogroups with species gene abundances greater than 5.

### Diversity and evenness

*C. perspicillata* demonstrated the most diverse *OR* repertoire among the three species (Fig. 2A; *H*’=6.33; *t*_*perspicillata*_ (16)=2.8, *p*_*perspicillata*_ = 0.013) and *C. castanea* had the least diverse (H’=6.06), while *C. sowelli* had a diversity that fell within the middle (*H*’=6.22; *t*_*sowelli*_(16)=1.7, *p*_*sowelli*_ = 0.104; Fig. 2A). No apparent differences in evenness were observed (Fig. 2A; *t*_*sowelli*_(16)= −0.58; *p*_*sowelli*_=0.57; *t*_*perspicillata*_(16)= −1.13; *p*_*perspicillata*_=0.28;). Among *OR* subfamilies (Fig. 2B), *C. perspicillata* also consistently had the most diverse and *C. castanea* the least diverse, with the exception of OR56 (*C. sowelli* most diverse) and OR11 (*C. sowelli* and *C. castanea* both more diverse). Evenness among OR subfamilies was similar across all species (Fig. 2B), though OR11 and OR1/3/7 show less evenness in *C. sowelli* and OR4 shows much less evenness in *C. castanea* relative to the other species.

## Discussion

Ecological specialization is expected to be linked to trait diversity, with generalist species exhibiting traits that enable access to a wider range of resources. We tested this hypothesis using three species of closely, related neotropical short-tailed fruit bats (*Carollia*) with overlapping geographic ranges, but a spectrum of dietary specialization on *Piper* fruits. We applied a new approach, ecological diversity indices, to examine how the *OR*s of these bats varies with increasing ecological specialization.

Measuring diversity among orthogroups provides deeper evolutionary insight than simply comparing numbers of genes and may illuminate the evolutionary processes and functions underlying current diversity. For example, *C. perspicillata* technically has more *OR*s in subfamily OR5/8/9 (Fig. 1B), but measures of diversity and evenness are actually quite similar across the three species (Fig. 2B). In contrast, subfamily OR1/3/7 shows substantial differences in diversity among the three species (Fig. 2B) while the receptor counts within this subfamily are difficult to interpret (Fig. 1B). *OR*s are among the fastest evolving genes in the genome (Yohe et al. 2020b), and their turnover via birth-death evolution makes it challenging to compare orthologs among species. For example, there have been so many *OR* gains and losses within rodents that there is less 70% homology in *OR*s and less than 20% homology in *vomeronasal type-1* genes (another chemoreceptor gene family) between a mouse and a rat (Zhang et al. 2007). The number of receptors only becomes meaningful in terms of describing the “diversity” of receptors in the repertoire, and increased numbers of orthogroups may indicate more potential ligands to be perceived. Thus, if a species has more orthogroups, then there are more distinct forms of *OR*s present, and subsequent paralogs within these orthogroups reinforce the diversity. However, fewer orthogroups and increased paralogs suggest redundancy within an orthogroup. This increased redundancy may suggest selection for retention of similar paralogs, and potentially have a favorable dosage effect (Teufel et al. 2016; Yohe et al. 2019b). Tandem gene duplicates are often expressed even greater than twofold, with dramatically higher activity than other sites in the genome (Loehlin and Carroll 2016). Even if increased dosage of expression is not observed, selection for duplicate retention and increased redundancy may also be advantageous if the receptor is critical to detecting a food resource. Olfactory sensory neurons stochastically express a single *OR* gene (Rodriguez 2013; Monahan and Lomvardas 2015), and multiple tandem copies of a gene of similar function may increase the probability of expression. In other words, having multiple copies of a similar receptor may increase its chances of expression. Alternatively, more paralogs may indicate divergent function. While counterintuitive, functional evidence in primates suggests that orthologous *OR*s across divergent species are more likely to bind to the same odorant ligand than paralogs (Adipietro et al. 2012). Given the low levels of codon substitution observed in our gene trees, however, we predict that paralogs might be more similar in function and advocate for the dosage effect hypothesis in *Carollia*.

We found that the more generalist frugivorous bat, *C. perspicillata*, has a more diverse collection of distinct *OR*s compared to the specialist *C. castanea*. The interpretation of our results relies on the assumption that an increased number of different orthogroups (not number of intact genes) reflects an increased potential to detect different odorant ligands. For example, during the transition from specialist to generalist diet in nymphalid butterflies (*Vanessa*), the generalist species expanded their gustatory receptor repertoire and this increased repertoire size is associated with a more diverse plant resource use (Suzuki et al. 2018). However, instead of measuring increased gene birth rates, we measure the result of gene duplicate retention as a function of diversity of different receptors in the genome. While the former assumes that duplication rates are deterministic and not stochastic processes, the latter, focused on diversity within orthogroups may more correctly reflect products of selection. In *Carollia*, because more than 50% of the diet of *C. perspicillata* relies on a diversity of plant resources other than *Piper* (Fig. 1A; e.g., Fleming 1991; Maynard et al. 2019), the number of different compounds this species needs to detect should be greater than that of the species that primarily consume fruits within the *Piper* genus. Given the overlapping geographic distributions and dietary niches, divergent olfactory profiles among these *Carollia* species may optimize for the detection of different plant resources in a cluttered rainforest community.

While which odorant ligands bind to which *OR*s in bats is completely unknown, our analyses constitute a major contribution to help isolate clusters of receptors as candidates to functionally investigate whether relevant environmental scent cues initiate a response for these receptors. Because total numbers of intact receptors are irrelevant to olfactory function, exceptional retention of recent gene duplicates and orthogroups containing overrepresentation of species-specific in-paralogs may be a more meaningful starting point for deorphanization. With this approach, instead of attempting to decode hundreds of receptors, 10-20 genes become good experimental candidates. For example, the *Piper* specialist *C. castanea* shows behavioral preference and attraction to volatile cues of ripened *P. sancti-felicis* fruits (Maynard et al. 2019; Leiser-Miller et al. 2020). Thus, a future study may test the hypothesis whether receptors demonstrating exceptional redundancy within *C. castanea* (e.g., such as those found in OR4 (Fig. 2B) or OR10 (Fig. 3)) respond to volatiles of ripened fruits of *P. sancti-felicis* in a biochemical assay. Detecting olfactory adaptation at the molecular level in olfaction remains an open challenge (Yohe and Brand 2018), yet our discovery of inverse patterns of dietary specialization and *OR* diversity may have consequential implications for understanding how evolution shapes complex and rapidly-evolving gene families.

**Figure 3.**
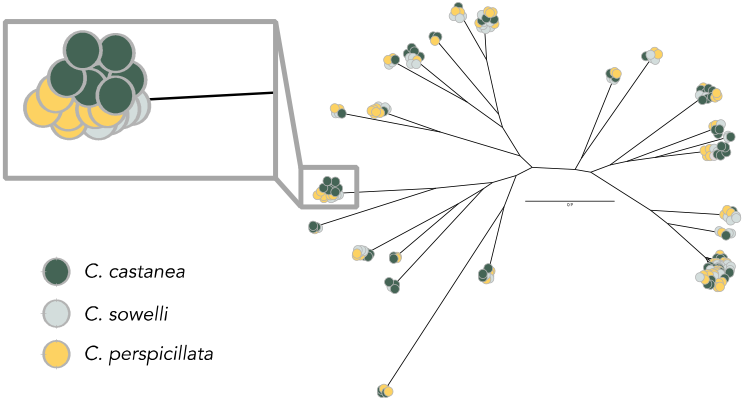
Inferred codon-model gene tree for *olfactory receptor* subfamily 10 (*OR*10). Each colored circle represents an *OR* gene colored by species. Larger clusters of genes are orthogroups or clusters of orthogroups that include orthologous genes and paralogs. The window inset indicates a cluster of related genes indicated as exceptional representation in Figure 2 for *C. castanea*.

## Acknowledgments

We thank the faculty and staff of La Selva Biological Research Station in Costa Rica for hosting our research team. For support during fieldwork, we thank K.T. Davies and S.J. Rossiter, and for support with paperwork, we thank J. Hurtado and C.M. Orrego. For illustrations, we thank C.M. Mauro. This project was funded through support from the National Science Foundation (NSF) Graduate Research Fellowship, Society for the Study of Evolution Rosemary Grant, American Society of Mammalogists, NSF-PRFB 1812035, and NSF-IOS 2032073 to LRY; NSF-DEB 1701414 to LRY and LMD; NSF-DEB1442142, NSF-DEB 1456455 and NSF-IOS 2031906 to LMD; NSF-DEB 1456375 to SES. The authors declare no conflicts of interest.

## Author Contributions

LRY designed the study, aided in collecting the samples, sequenced the genes, ran analyses, and wrote the manuscript. LMD designed the study; LMD and SES edited the manuscript.

## Data Accessibility

Raw Illumina sequence reads from targeted sequence capture were deposited to GenBank (SRAXXXX). Alignments, gene trees, and orthogroup results were deposited into Dryad (DOI: XXXXXX).

