## supplement for "Diversity in olfactory receptor repertoires is associated with dietary specialization in a genus of frugivorous bat"

**Table S1**. Field information for specimens used in this study.

| **Species** | **Field Number** | **Locality** | **Date Captured** | **Sex** |
| --- | --- | --- | --- | --- |
| *Carollia castanea* | LS084 | La Selva, Costa Rica | Aug 4 2017 | M |
| *Carollia sowelli* | LS073 | La Selva, Costa Rica | Aug 4 2017 | M |
| *Carollia perspicillata* | LS070 | La Selva, Costa Rica | Aug 4 2017 | M |

**Table S2**. Number of intact olfactory receptor (OR) genes found within each olfactory receptor subfamily. Intact OR genes are genes with reading frames longer than 650bp that do not have a premature stop codon or frameshift mutation.

| **Species** | **Class I** | | | | **Class II** | | | | | | | | |  |
| --- | --- | --- | --- | --- | --- | --- | --- | --- | --- | --- | --- | --- | --- | --- |
|  | **51** | **52** | **55** | **56** | **1/3/7** | **2/13** | **4** | **5/8/9** | **6** | **10** | **11** | **12** | **14** | **Total** |
| *Carollia castanea* | 73 | 69 | 1 | 16 | 188 | 102 | 103 | 180 | 56 | 74 | 14 | 3 | 2 | 881 |
| *Carollia sowelli* | 75 | 69 | 1 | 24 | 261 | 127 | 99 | 200 | 62 | 73 | 22 | 1 | 2 | 1017 |
| *Carollia perspicillata* | 93 | 104 | 3 | 15 | 258 | 121 | 98 | 242 | 83 | 76 | 14 | 4 | 4 | 1115 |
| Orthogroup count | 75 | 84 | 1 | 13 | 236 | 117 | 116 | 210 | 66 | 77 | 19 | 3 | 3 | 1019 |

**Table S3**. Number of intact sequences for each olfactory receptor subfamily. Alignment length and percent identity was calculated from nucleotide alignments based on the transAlign algorithm. The model of evolution estimated for each alignment was determined by ModelOMatic v.1.01 and applied to each alignment for tree inference in IQtree. Data in this table refer to the alignment with the root, but with stop codons for all sequences removed.

| **OR subfamily** | **# sequences** | **Alignment Length (bp)** | **% pairwise identity** | **Model** |
| --- | --- | --- | --- | --- |
| *Class I* |  |  |  |  |
| OR 51 | 242 | 1,149 | 58.5 | Codon F64+4dG |
| OR 52 | 243 | 1,212 | 58.5 | Codon F64+4dG |
| OR 55 | 7 | 1,074 | 80.2 | Codon F64+4dG |
| OR 56 | 56 | 1,095 | 79.2 | Codon F64+4dG |
| *Class II* |  |  |  |  |
| OR 1/3/7 | 708 | 1,242 | 64.9 | Codon F3X4+4dG |
| OR 2/13 | 351 | 1,191 | 57.8 | Codon F1X4+4dG |
| OR 4 | 301 | 1,263 | 63.0 | Codon F64+4dG |
| OR 5/8/9 | 623 | 1,284 | 57.5 | Codon F1X4+4dG |
| OR 6 | 202 | 1,194 | 59.6 | Codon F64+4dG |
| OR 10 | 224 | 1,167 | 55.1 | Codon F64+4dG |
| OR 11 | 51 | 1,128 | 64.8 | Codon F64+4dG |
| OR 12 | 9 | 1,065 | 84.7 | Codon F3X4+4dG |
| OR 14 | 9 | 1,104 | 77.9 | Codon F3X4+4dG |
